# Evidence for the coupling of substrate recognition with transporter opening in MOP-family flippases

**DOI:** 10.1101/260661

**Authors:** Lok To Sham, Sanduo Zheng, Andrew C. Kruse, Thomas G. Bernhardt

**Affiliations:** Department of Microbiology and Immunobiology, Harvard Medical School, Boston, MA 02115; Department of Biological Chemistry and Molecular Pharmacology, Harvard Medical School, Boston, MA 02115; Department of Microbiology and Immunology, National University of Singapore, Singapore, 117545

**Keywords:** peptidoglycan, cell wall, MurJ, flippase, MOP-family transporters

## Abstract

Bacteria produce a variety of surface-exposed polysaccharides important for cell integrity, biofilm formation, and evasion of the host immune response. Synthesis of these polymers often involves the assembly of monomer oligosaccharide units on the lipid carrier undecaprenyl-phosphate at the inner face of the cytoplasmic membrane. For many polymers, including cell wall peptidoglycan, the lipid-linked precursors must be transported across the membrane by flippases to facilitate polymerization at the membrane surface. Flippase activity for this class of polysaccharides is most often attributed to MOP (Multidrug/Oligosaccharidyllipid/Polysaccharide) family proteins. Little is known about how this ubiquitous class of transporters identifies and translocates its cognate precursor over the many different types of lipid-linked oligosaccharides produced by a given bacterial cell. To investigate the specificity determinants of MOP proteins, we selected for variants of the WzxC flippase involved in *Escherichia coli* capsule (colanic acid) synthesis that can substitute for the essential MurJ MOP-family protein and promote transport of cell wall peptidoglycan precursors. Variants with substitutions predicted to destabilize the inward-open conformation of WzxC lost substrate specificity and supported both capsule and peptidoglycan synthesis. Our results thus suggest that specific substrate recognition by a MOP transporter normally destabilizes the inward-open state, promoting transition to the outward-open conformation and concomitant substrate translocation. Furthermore, the ability of WzxC variants to suppress MurJ inactivation provides strong support for the designation of MurJ as the flippase for peptidoglycan precursors, the identity of which has been controversial.

**SIGNIFICANCE:** From cell walls in bacteria to protein glycosylation in eukaryotes, surface exposed polysaccharides are built on polyprenol-phosphate lipid carriers. Monomer units are typically assembled at the cytoplasmic face of the membrane and require translocation to the cell surface for polymerization/assembly. MOP-family proteins are a major class of transporters associated with this flippase activity. Despite their ubiquity and importance for cell surface biology, little is known about their transport mechanism. Here, we investigated substrate recognition by MOP transporters in bacteria. We present evidence that transport proceeds via destabilization of the inward-open state of the transporter by specific substrate binding thereby promoting a transition to the outward-open state and substrate release on the opposite face of the membrane.

## INTRODUCTION

Bacterial cells produce a variety of cell surface polysaccharides. Polymers like peptidoglycan (PG) and teichoic acids (TAs) are critical for cell integrity and shape maintenance, whereas capsular polysaccharides and O-antigens play central roles in virulence and the evasion of host defenses (1). Despite their vast structural diversity, the majority of surface polysaccharides are made by one of three types of synthesis and export mechanisms: synthase-dependent, ABC transporter-dependent, or Wzy-dependent pathways (1). Most complex polysaccharides with 3-6 sugars in their repeating unit are made by the widely-distributed Wzy-dependent pathway (2). Examples include O-antigens of gram-negative bacteria and many capsular polysaccharides. For these polymers, the monomeric oligosaccharide unit is synthesized at the inner face of the cytoplasmic membrane on the lipid carrier undecaprenyl phosphate (Und-P) (2, 3). For polymerization, the oligosaccharide moiety must be transported across the membrane and exposed at the membrane surface by a class of transporters referred to as flippases (3, 4). Notably, this overall lipid-linked synthesis strategy is near universal as it is also employed by eukaryotic cells to generate oligosaccharides for N-linked protein glycosylation (5).

Lipid-linked sugar flippase activity for polysaccharide synthesis has principally been ascribed to one of two types of transporters: ABC (ATP-binding cassette) systems, and MOP (Multidrug/Oligosaccharidyl-lipid/Polysaccharide) family proteins (3). MOP-type flippases are typically associated with Wzy-dependent polysaccharide synthesis pathways, and several lines of evidence indicate that they have a strong preference for translocating the specific lipid-linked oligosaccharide produced by their cognate synthetic pathway (6–8). How these transporters specifically recognize their substrates over the many different types of lipid-linked oligosaccharides produced in a bacterial cell is not currently known. The transport mechanism is also poorly understood, but the recently solved structures of the MOP-family protein MurJ from *Thermosipho africanus* and *Escherichia coli* has provided important clues (9, 10).

MurJ was identified several years ago as a protein essential for PG biogenesis. It was proposed to be the long-sought after flippase that translocates the final PG precursor lipid II (11, 12), which consists of the disaccharide N-acetylmuramic acid (MurNAc)-β-1-4-N-acetylglucosamine (GlcNAc) with a pentapeptide attached to the MurNAc sugar via a lactyl group. Like other polysaccharide synthesis pathways, lipid II is assembled on the inner face of the cytoplasmic membrane and must be translocated before it can be polymerized and crosslinked to form the PG matrix that fortifies bacterial cells against osmotic rupture (3). Importantly, the functional assignment of MurJ as the PG lipid II flippase remains controversial in the field of PG biogenesis because in vitro assays have detected what appears to be lipid II flippase activity for the SEDS (shape, elongation, division, sporulation) family protein FtsW (3, 13).

In support of a flippase function for MurJ, it was found to be required for lipid II translocation in *Escherichia coli* using an *in vivo* flippase assay whereas SEDS proteins were dispensable for this activity (12). The structure of MurJ from *T. africanus* also supports a transporter function (9). In the crystals, the protein adopted an inward-open conformation with a solvent accessible cavity capable of accommodating the lipid II head group. Related structures of a different subfamily of MOP transporters involved in drug efflux had previously been solved in an outward-open conformation (14), and chemical probing of MurJ structure in vivo (15) indicates that MurJ can adopt such a conformation in cells. Evolutionary coupling (EC) analysis (16) also shows that MurJ must adopt an additional conformation not accounted for in the inward-open crystal structure, since pairs of distant residues on the cytosolic face so strong co-evolution. An outward-open model of MurJ similarly leaves predicted interactions unsatisfied on the periplasmic face of the protein, indicating that at least two distinct conformations are subject to evolutionary selective pressure (10). Thus, it has been proposed that MurJ and other MOPS family proteins mediate transport/flipping via an alternating access model involving the interconversion between the inward- and outward-facing conformations, alternately allowing the lipid II headgroup to access the cytosolic and periplasmic faces of the membrane (2, 3, 9, 10).

To investigate the specificity determinants of MOP proteins, we selected for variants of the WzxC flippase involved in *Escherichia coli* capsule (colanic acid) synthesis (17) that gain translocation activity for peptidoglycan precursors and can substitute for the essential MurJ MOP-family protein in the cell wall synthesis pathway. Variants with amino acid changes predicted to destabilize the inward-open conformation of WzxC lost substrate specificity and supported both capsule and peptidoglycan synthesis. Our results thus suggest that specific substrate recognition by a MOP transporter normally functions to destabilize the inward-open state, promoting transition to the outward-open conformation and concomitant substrate translocation. Furthermore, the ability of WzxC variants to suppress MurJ inactivation provides strong support for the designation of MurJ as the flippase for peptidoglycan precursors.

## RESULTS

### Identification of WzxC variants that can substitute for MurJ

WzxC is a MOP-family transporter in *E. coli* required for the synthesis of the colanic acid capsule (17), the production of which is induced by activation of the Rcs envelope stress response (18). The colanic acid precursor is a hexasaccharide [L-fucose-(pyruyl-D-galactose-D-glucouronic acid-D-galactose)-O-acetyl-L-fucose-D-glucose] built on the Und-P lipid carrier. WzxC has been implicated in the transport (flipping) of this lipid-linked intermediate (17). Although WzxC is the closest relative of MurJ in *E. coli* (≈12% sequence identity), its substrate structure differs greatly from the lipid II PG precursor that MurJ has been implicated in flipping. It is therefore not surprising that WzxC fails to substitute for MurJ and promote growth when MurJ is depleted (**Fig. 1, row 1**). However, we thought it might be possible to identify altered WzxC proteins that gain the ability to flip the PG precursor and rescue a MurJ defect. We reasoned that the isolation and characterization of such variants would provide useful information about what determines substrate specificity in MOP-family flippases and potentially reveal new insights into the transport mechanism.

**Figure 1.**
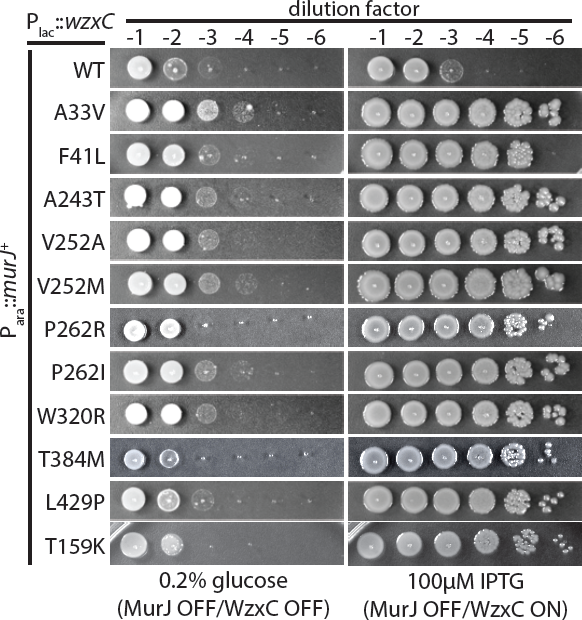
WzxC variants can substitute for MurJ. Cells of CS7 [P*_ara_*::*murJ*] harboring plasmids encoding C-terminally FLAG-tagged WzxC (WT) or the indicated derivatives were grown in LB medium with arabinose overnight. Following normalization for culture density, serial dilutions (10^-1^ to 10^-6^) were prepared and 5 µl of each were spotted onto LB plates supplemented with either glucose or IPTG. Plates were photographed after incubation at 37^o^C for ≈16 hours. All of the strains grew similarly on plates supplemented with arabinose under this condition. An additional growth experiment with additional concentrations of IPTG are presented in **SI Appendix, Fig. S2.**

To select for altered substrate specificity variants of WzxC, the *wzxC* gene was mutagenized using error-prone PCR and cloned into a medium copy vector under control of the lactose promoter (P*_lac_*). The resulting plasmid library was transformed into a MurJ-depletion strain where native *murJ* was engineered to be under control of the arabinose promoter (P*_ara_*). This promoter replacement renders the strain dependent on the presence of arabinose in the medium for growth. When the depletion strain harboring the *wzxC* plasmid library was plated on LB medium lacking arabinose but supplemented with isopropyl-β-D-thiogalactopyranoside (IPTG) to induce *wzxC*, surviving colonies arose at a low frequency (10^-4^). To distinguish between survivors with mutations allowing arabinose-independent expression of *murJ* from the desired *wzxC* mutants capable of substituting for *murJ*, plasmids were purified from the isolates and transformed back into the parental MurJ-depletion strain. The resulting transformants were then tested for growth on IPTG-containing medium with or without arabinose supplementation. Plasmids conferring arabinose-independent growth were then isolated and their *wzxC* insert was sequenced. Many of the primary isolates harbored *wzxC* clones with multiple mutations (**SI Appendix, Table S1**). To identify the functionally relevant substitutions, we used site-directed mutagenesis to construct plasmids encoding C-terminal FLAG-tagged WzxC variants (WzxC-FLAG) with single amino acid changes corresponding to those identified in the original mutant isolates. The FLAG-tag did not appear to interfere with WzxC activity as the fusion was capable of supporting capsule production in a Δ*wzxC* strain (**SI Appendix, Fig. S1**). Importantly, wild-type WzxC-FLAG also failed to promote growth upon MurJ depletion (**Fig. 1, row 1**). In total, eleven single amino acid substitutions in WzxC-FLAG representing changes throughout the length of the 492 amino acid protein were found to be sufficient for suppression of MurJ depletion (**Fig. 1, rows 2-11**). The *wzxC* alleles varied in the strength of the observed suppression phenotype, with most being able to promote growth of the MurJ depletion strain upon induction with 25 µM IPTG, but two requiring 75-100 µM IPTG to achieve full suppression (**Fig. 1, rows 2-11, SI Appendix, Fig. S2**). Immunoblot analysis using anti-FLAG antibodies indicated that all of the WzxC-FLAG variants were produced at levels comparable to, or slightly lower than, the wild-type protein (**SI Appendix, Fig. S3**). Thus, the WzxC variants do not gain the ability to substitute for MurJ simply due to their overproduction.

To determine if the altered WzxC proteins could fully substitute for MurJ, we assessed the ability of eight variants for their ability to support the growth of a *murJ* deletion. A Δ*murJ*::Kan^R^ allele constructed in a background with a complementing *murJ* plasmid was used as a donor for P1 phage transduction of the deletion into MurJ^+^ strains harboring the *wzxC* plasmids. For all the mutants tested, transductants were successfully isolated on medium containing IPTG for *wzxC* induction, and deletion/replacement of the native *murJ* gene was confirmed in each case (**SI Appendix, Fig. S4**). We conclude that many of the WzxC variants identified in the selection are capable of overcoming the complete loss of MurJ function. Therefore, we will henceforth refer to them as ^MJ^WzxC derivatives.

### ^MJ^WzxC variants can support PG lipid II flipping in vivo

The ability of the ^MJ^WzxC variants to suppress the essentiality of MurJ suggests that they have gained the ability to transport the lipid II precursor for PG biogenesis. To test this possibility, we took advantage of an in vivo assay for lipid II flipping (12). To detect lipid II transport, cells were radiolabeled with the PG precursor [^3^H]-meso-diaminopimelic acid (mDAP) and treated with Colicin M (ColM). This toxin invades the periplasm and cleaves flipped lipid II, generating a soluble pyrophospho-disaccharide pentapeptide that is subsequently converted to disaccharide tetrapeptide by periplasmic carboxypeptidases. When MurJ is functional, ColM cleavage of flipped lipid II generates a new soluble radiolabeled product and destroys the labeled lipid fraction (12, 19) (**Fig. 2**). The MurJ variant, MurJ(A29C), is sensitive to the Cys-modifying reagent (2-sulfonatoethyl)methanethiosulfanate (MTSES). When radiolabeled cells relying on this MurJ derivative for lipid II translocation are treated with MTSES, the lipid fraction is protected from ColM cleavage and the soluble ColM product is not observed (12) (**Fig. 2**), indicating that flipping is blocked. A plasmid producing WzxC(WT) did not alleviate this block (**Fig. 2**). However, production of WzxC(V252M) restored lipid II cleavage by ColM in MurJ-inactivated cells and promoted the accumulation of the soluble ColM product (**Fig. 2**). We therefore conclude that the ^MJ^WzxC derivatives have gained the ability to facilitate lipid II transport.

**Figure 2.**
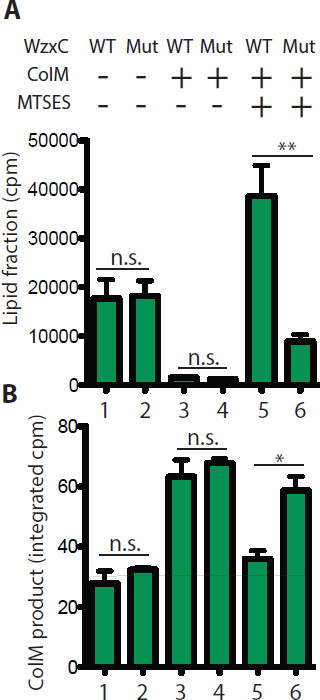
Support of PG lipid-II flipping in vivo by a WzxC variant. Cells of CAM290 [*murJ*(A29C)] harboring plasmid pCS124 [P*_lac_*::*wzxC*(WT)-*flag*] or pDF2 [P*_lac_*::*wzxC*(V252M)-*flag*] were grown in labeling medium to an OD_600_ of 0.2. [^3^H]-mDAP was then added to radiolabel PG precursors. After a 15 min labeling period, MTSES and ColM were added to block MurJ(A29C) activity and cleave flipped lipid II, respectively. Just prior to cell lysis, cells were collected by centrifugation and fractionated to measure radioactivity in the PG lipid precursor pool (**A**) and soluble ColM cleavage product (**B**). WT and Mut denote WzxC(WT) and WzxC(V252M), respectively. The means and the standard error of means (SEMs) from three experiments are shown. P-values were calculated with two-tailed unpaired Student’s t-test. *, p < 0.05; **, p < 0.01; n.s., not significant. cpm = counts per minute.

### Most ^MJ^WzxC derivatives have lost substrate specificity

We next investigated whether the ^MJ^WzxC variants that support lipid II translocation retain the ability to promote colanic acid capsule synthesis. Production of the capsule is induced by the Rcs stress response system when cells are grown in high-salt medium (20). Cells unable to make capsule grew poorly on LB medium containing 1.0 M NaCl (**Fig. 3A, row 1, SI Appendix, Fig. S1**). This growth defect was observed for mutants blocked at the first step in the pathway (Δ*wcaJ*) or at the WzxC step. Thus, the phenotype does not require the build up of lipid-linked colanic acid precursors that would likely reduce the pool of lipid carrier available to other pathways like PG synthesis (21). Production of wild-type WzxC-FLAG as well as most of the ^MJ^WzxC-FLAG derivatives restored the growth of Δ*wzxC* cells on LB with 1.0M NaCl (**Fig. 3A, rows 2-13**). The exception was WzxC(T159K), which suppressed MurJ depletion, but not the Δ*wzxC* phenotype. Thus, the majority of the ^MJ^WzxC proteins appear to retain WzxC function.

**Figure 3.**
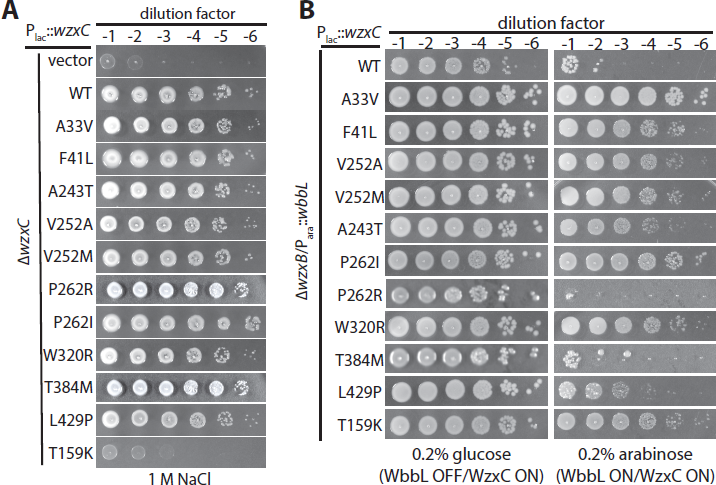
Transport of colanic acid and O-antigen precursors by ^MJ^WzxC variants. (A) Cells of CS38 [∆*wzxC*] harboring an empty vector (vector) or vectors encoding the indicated FLAG-tagged WzxC variant were grown and plated on LB medium with 1M NaCl as described in Figure 1. Complementation of the ∆*wzxC* phenotype results in the formation of mucoid colonies on the high-salt medium. **(B)** Cells of CS39/pCS160 [∆*wzxB*/P*_ara_*::*wbbL*] harboring the same WzxC-encoding plasmids were grown and diluted as descried in Figure 1 followed by plating on LB medium with 100 µM IPTG (to induce WzxC production) plus either glucose or arabinose (to repress or induce O-antigen production, respectively) as indicated.

To further investigate the range of potential substrates capable of being utilized by the ^MJ^WzxC variants, we tested their ability to participate in O-antigen synthesis. These polysaccharides decorate the lipopolysaccharides (LPS) that form the outer leaflet of the outer membrane in gram-negative bacteria (22). *E. coli* K-12 strains, including the MG1655 derivatives used here, do not make O-antigen polymers. This defect is due to an insertion element that disrupts *wbbL*, which encodes the enzyme catalyzing the committed step for the synthesis of the O-16 antigen with the repeating unit D-galactose-D-glucose-L-rhamnose-(D-glucose)-D-N-acetylglucosamine (23). The flippase for this pathway is thought to be WzxB (24). Inactivation of *wzxB* is not lethal in WbbL^-^ strains. However, ectopic expression of *wbbL* in Δ*wzxB* cells is lethal, presumably due to the accumulation of lipid-linked O-16 precursors and the sequestration of Und-P lipid carrier from the PG synthesis pathway (21, 25). Unlike wild-type WzxC-FLAG, co-production of many of the ^MJ^WzxC-FLAG variants wth WbbL rescued the WbbL-induced lethality of Δ*wzxB* cells and restore O-antigen production (**Fig. 3B, SI Appendix, Fig. S5**). However, two variants, WzxC(P262R) and WzxC(T384M) that complemented the MurJ-depletion and Δ*wzxC* phenotypes failed substitute for WzxB. On the other hand, WzxC(T159K), which failed to complement Δ*wzxC* rescued both the Δ*wzxB* and MurJ-depletion phenotypes. From these results, we conclude that the ^MJ^WzxC variants have largely lost substrate specificity, allowing them to function in a variety of polysaccharide synthesis pathways.

### Substitutions in the ^MJ^WzxC variants are predicted to destabilize the inward-open conformation

In order to better understand the molecular basis for the effects of the ^MJ^WzxC mutations, we constructed a homology model of WzxC using the crystal structure of *E. coli* MurJ (10) as a template (**Fig. 4, SI Appendix, Fig. S5**). We expected that most of the specificity altering changes in WzxC would occur in the aqueous cavity of the transporter where substrate is predicted to bind. However, when mapped onto the model WzxC structure, the majority of residues altered in the ^MJ^WzxC variants clustered at or near the periplasmic face of the protein. Many of these substitutions occur in residues that mediate inter-domain contact between the N-lobe and C-lobe on the periplasmic side (A33, A243, F41 and W320). For example, the pseudo-symmetry related pair of residues A33 and A243 are in small, sterically restricted spaces (**SI Appendix, Fig. S5A**). Mutation of these to Val or Thr is incompatible with the inward-open homology model due to steric clash with neighboring residues, and so is expected to destabilize the inward-open conformation. Similarly, the buried N-lobe residue F41 engages in extensive hydrophobic contacts with L248, F316, and other residues in the C-lobe, so its mutagenic substitution with Leu may destabilize the inward-open conformation (**SI Appendix, Fig. S5A**). Likewise, W320 sits near the interface between the two lobes, and the nonconservative substitution with Arg likely weakens interdomain interactions on the periplasmic side (**SI Appendix, Fig. S5A**). It is noteworthy that most of these substitutions are modest. They may simply raise the energy of the inward-open state, altering the equilibrium between inward-open and outward-open states without causing a complete loss of function.

**Figure 4.**
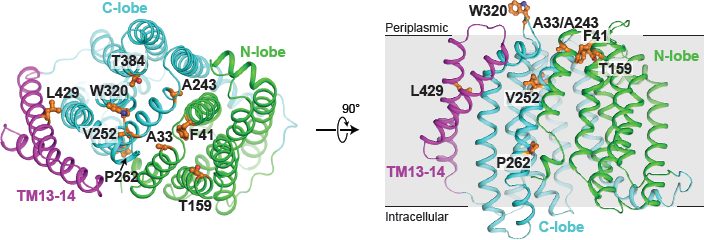
Structural analysis of specificity-broadening mutations in WzxC. A homology model of WzxC is shown, with the sites of mutations highlighted in orange sticks. At left, the protein is viewed from the periplasmic side, and at right is viewed parallel to the membrane plane. With only a few exceptions, the mutations cluster at the periplasmic face of the protein near the interface between the N- and C-terminal lobes of WzxC.

Other substitutions are not located at the interdomain interface but rather are found in or near the central cavity of the enzyme. For instance, P262 and V252 sit in the lateral gate between TM1 and TM8 (**SI Appendix, Fig. S5B**). Both residues may be directly involved in substrate binding or play a critical role in conformation transition. Their alteration may expand the range of substrates accepted by transporter by affecting substrate binding affinity. Notably, L429 is located neither in the central cavity nor in the extracellular gate, but rather is found in TM13 (**SI Appendix, Fig. S5C**). Its mutation to proline is incompatible with α-helical geometry, and must force a distortion in the helix. The connection between this effect and broadened substrate specificity is unclear, but attests to a functionally important role for TMs 13 and 14, which are absent in most MOP family flippases.

## DISCUSSION

In this report, we isolated ^MJ^WzxC variants that have lost substrate specificity and gain the ability to transport the PG precursor lipid II and O-antigen precursors in addition to its native substrate for colanic acid synthesis. The location of the amino acid changes in WzxC resulting in this phenotype was surprising. Rather that altering the predicted substrate-binding region of the modeled structure, the changes largely map to portions of the protein located near the periplasmic face of the membrane. Based on the MurJ structure, this region of the protein is predicted to form contacts that stabilize the inward-open conformation of the transporter. Because many of the amino acid substitutions in the ^MJ^WzxC variants involve a change in side chain size or charge, we infer that they exert their effect on the transporter through the destabilization of the inward-open conformation. Thus, the genetic results support a model in which the stability of the inward-open conformation plays a key role in determining the substrate specificity of the transporter (**Fig. 5**).

**Figure 5.**
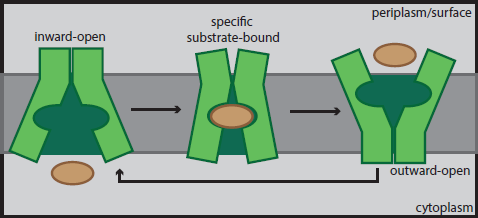
Model for substrate induced conformational changes in MOP family flippases. Shown is a schematic summarizing our model for substrate transport by MOP family flippases. Structural studies suggest that the inward-open conformation is the most stable state of the transporter. Based on our genetic results, we propose that specific substrate binding is required to break contacts at the outer face of the transporter to destabilize the inward-open conformation. Once these contacts are broken, a transition to the outward-open conformation can occur to allow for substrate release on the opposite face of the membrane. For simplicity, the lipid anchor of the substrate is not drawn.

We propose that for wild-type WzxC, a specific interaction between the native substrate and the hydrophillic core of the transporter is needed to break contacts at the periplasmic face of the membrane involved in stabilizing the inward-open conformation. Such a change would then facilitate the transition to the outward-open conformation and the release of substrate on the outside face of the membrane. Once substrate is released, the protein would then be free to transition back to the more stable inward-open conformation and begin another round of transport. Due to the changes in the ^MJ^WzxC variants, we envision that the protein can more readily interconvert between the two main conformations without the need for specific substrate binding. Thus, the transport of non-native precursors would be facilitated by the altered flippase. The main limitation in this case is likely to be the ability of the precursor sugar moiety to fit within the hydrophillic core of the altered transporter. WzxC was therefore an especially fortuitous choice for this specificity study. Its native substrate is relatively large such that the hydrophillic core of the ^MJ^WzxC variants is likely capable of accommodating and flipping a wider range of substrates than might be possible with other transporters.

The phenotype of a previously isolated mutant in a different flippase suggests that other transporters may function similarly to WzxC. TacF is a MOP family protein implicated in the transport of teichoic acid (TA) precursors of *Streptococcus pneumoniae* (26). The LTAs in this organisms are normally decorated with choline (27). A TacF variant was identified that suppressed the choline-dependent growth phenotype of *S. pneumoniae*, presumably by allowing the transport of LTA precursors lacking choline (26). The change in this variant that alters the substrate choline requirement is located in a loop of TacF predicted to be at the outer surface of the membrane (26). Similar to WzxC, this area is exactly where contacts that stabilize the inward-open conformation are likely to be made. Thus, the use of a specific substrate binding event to destabilize the inward-open state and promote a conformational transition may be a general component of the transport mechanism of MOP family flippases. Substrate-induced conformational changes have also been implicated in the transport mechanism of the (NSS) family of transporters (28), suggesting that they may be involved in many different types of membrane transport processes.

In addition to a better understanding of the transport mechanism of MOP family flippases, the activities of the ^MJ^WzxC variants also provide insight into the process of PG biogenesis. Although flippase activity has yet to be demonstrated for MurJ in vitro, the finding that a protein implicated in flipping colanic acid precursors can substitute for MurJ in PG biogenesis makes it hard to argue that MurJ is anything other that the lipid II flippase. Furthermore, the ability of MJWzxC variants as well as other heterologous or promiscuous flippase proteins to maintain growth and viability upon MurJ inactivation suggests that lipid II transport does not need to occur in the context of specific multi-protein complexes with other PG biogenesis factors. Such complexes may be formed to render the process more efficient, but they do not appear to be necessary for the construction of the PG layer.

In conclusion, our results highlight the utility of unbiased genetic selections to study the function of MOP family flippases. Further structural analysis of these transporters using the substitutions identified in the ^MJ^WzxC variants should facilitate the capture of additional conformations of these proteins and provide further insight into their transport mechanism.

## MATERIALS AND METHODS

### Media, Bacterial Strains and Plasmids

Strains used in this study are listed in **SI Appendix, Table S2**. Unless otherwise specified, *E. coli* cells were grown in lysogeny broth (LB) under aeration at 37^o^C. Where indicated, arabinose and glucose are added to a final concentrations of 0.2% (w/v). The antibiotics ampicillin, chloramphenicol, and kanamycin were used at a final concentration of 25 µg/ml. Spectinomycin was added to a final concentration of 40 µg/ml. Plasmids and oligonucleotides used in this study are listed in **SI Appendix, Table S3 and S4**, respectively.

### Selection of WzxC variants that can suppress MurJ essentiality

Strain CS7 [P_ara_::*murJ*] was transformed with a mutagenized *wzxC* plasmid library (P_lac_::*wzxC*) (see **SI appendix** for details). Transformants were scraped from the agar surface into 5ml of LB medium and the resulting cell suspension was serially diluted and plated on LB agar supplemented with chloramphenicol, 0.2% glucose, and IPTG. Isolates that required induction of the *wzxC* plasmid with IPTG for growth in the absence of *murJ* expression (0.2% glucose) were selected for sequencing.

### Detection of lipid II flippase activity using Colicin M

Lipid II translocation across the inner membrane was monitored using the previously described colicin M assay (12). Cells of CAM290/pCS124 [*murJ*(A29C)/P_lac_::*wzxC*] and CAM290/pDF2 [*murJ*(A29C)/P_lac_::*wzxC*(V252M)] were grown in LB medium with chloramphenicol overnight. Cultures were diluted 100 fold in 40ml of the labeling medium with 100µM IPTG (M9 supplemented with 0.1% (w/v) casamino acids, 0.2% (w/v) maltose, 0.1mg/ml of lysine, threonine and methionine) and grown at 37^o^C with aeration. When the culture OD_600_ reached 0.2, 15µl of 1.5 µCi/µl of ^3^H-mDAP (ARC) was added to 10ml of the culture and incubated for 15 minutes at 37^o^C. When indicated, colicin M and MTSES were added to a final concentration of 500 ng/ml and 0.4 mM, respectively. The cultures were then incubated for 10 minutes and chilled immediately on ice. Cells were collected by centrifugation at 8,000 x *g* for 2 minutes at 4^o^C and resuspended in 1ml of preheated water. Samples were then boiled for 30 minutes and processed to measure the soluble colicin M product and PG lipid precursors as described previously (12).

### Homology model construction

A homology model of *E. coli* WzxC was constructed in MODELLER (29) using a multi-template modeling protocol with the crystal structures of MurJ from *E. coli* (10) and *T. africanus* (9) (PDB ID: 5T77) serving as templates.

## ACKNOWLEDGEMENTS

The authors would like to thank all members of the Bernhardt, Rudner, and Kruse labs for helpful advice and discussions. This work was supported by the National Institutes of Health (R01AI083365 and AI099144 to TGB, and CETR U19 AI109764 to TGB and ACK).

